# Sex differences emerge after the menopause transition: Females show accelerated decline in episodic memory for spatial context at midlife

**DOI:** 10.1101/2024.05.06.591106

**Authors:** Annalise Aleta LaPlume, Rikki Lissaman, Julia Kearley, Maria Natasha Rajah

## Abstract

**Background and Objectives:** The ability to remember past events in rich contextual detail (episodic memory) declines with advancing age, with accelerated decline around midlife. Past research indicates there may be sex differences in cognitive aging trajectories and risk for age-related neurodegenerative diseases, i.e. Alzheimer’s Disease. Yet, little is known about how biological sex affects episodic memory in the adult lifespan. We examined age differences in episodic memory for spatial context in males and females.

**Research Design and Methods:** 192 adults aged 21 to 65 (*M*=44, *SD*=13, 134 females) completed a face-location task measuring spatial context memory (correct spatial context retrieval rates) and facial item memory (correct recognition rates), and the California Verbal Learning Test version II (CVLT-II) measuring verbal item memory (long free recall, cued recall, and recognition rates). Changepoint regression analysis was used to estimate the slope of memory across age and any significant shifts in the slope (indicating critical transition periods).

**Results:** Regression analyses revealed that the best-fitting model for females on spatial context memory accuracy was a one-changepoint model, with gradual decline of 2% (*SE*=1) fewer correct responses per year of age from age 21 until age 50 (95% CI 41, 58), shifting to more rapid decline of 4% (*SE*=1) fewer correct responses per year of age until age 65. The best fitting model for males on spatial context memory accuracy was linear, with no significant changes across ages. The best fitting models for both sexes were linear for facial item memory accuracy, spatial context memory and facial item memory reaction times, and verbal item memory accuracy.

**Discussion and Implication:** Males and females show similar decline on spatial context memory from young adulthood until midlife, after which females show greater decline than males. Importantly, disaggregating by sex indicated that past midlife effects on episodic memory for context may be driven by a specific group of females (post-menopausal), as accelerated decline occurred at the same time as menopause in midlife females and did not occur in midlife males.

**Translational Significance:** *Problem Addressed:* We use changepoint regression to examine how biological sex influences age differences in remembering the context of past events (episodic memory).

*Main outcome:* Females, but not males, showed significantly greater decline on spatial context memory (i.e., correctly recalling the location of previously learned information) after age 50, which aligns with the time of menopause in midlife females.

*Implications for Translation:* Episodic memory for spatial context shows accelerated decline in females after midlife compared to before midlife, but not in midlife males, indicating the potential influence of menopause on aging of memory.

## Background and Objectives

The ability to mentally re-experience an event or episode from a person’s past (e.g., your 21st birthday) is a form of long-term memory termed *episodic memory* (Tulving, 1972). Episodic memory is particularly sensitive to decline with advancing age (e.g., Fraundorf et al., 2019; Rönnlund et al., 2005; Park et al., 2002; Salthouse, 2002; Titz & Verhaeghen, 2010; Verhaeghen & Marcoen, 1993). Hence studying age-related variability in episodic memory is useful to the study of aging.

Advanced age particularly brings difficulties in remembering details from the circumstances of an event (e.g., the location of your 21st birthday) compared to other elements of the event, termed *context memory* (Spencer & Raz, 1995; Cansino et al., 2009; Rajah et al., 2010). The age-related difficulties with context memory are linked to an age-related difficulty on remembering associations between units of information for an event (termed *associative memory*) compared to remembering the individual units of information (*item memory*; Cansino, 2009; Naveh Benjamin, 2000, 2004; Old & Naveh-Benjamin, 2008). The difficulty in retrieving the spatial context of an item with age is also linked to age-related reductions in the ability to mentally re-experience an event (termed *recollection;* Jacoby & Kelley, 1992; Jacoby, 1991; Mandler, 1980; Yonelinas, 2002), rather than the dissociable process for having the impression or sense that an event has been previously experienced without retrieval of any additional information (termed *familiarity*; Koen & Yonelinas, 2014, 2016).

Several lifespan studies indicate age-related declines in context memory may occur at a non-linear rate. Some studies have found that age-related decline in context memory accelerates around the fifties and early sixties (Erngrund et al., 1996a, 1996b; LaPlume et al., 2022a; Uttl and Graf, 1993). Studies that specifically include a midlife group find that middle-aged and older adults have poorer retrieval accuracy than young adults on spatial and temporal context memory (Cansino et al, 2012; Ankudowich et al., 2016; Kwon et al., 2016). Notably, in one study mean retrieval accuracy did not differ between middle-aged and older adults, indicating that poorer context memory accuracy was present by middle age (Ankudowich et al., 2016). Middle-aged adults were also slower than young adults on retrieval reaction times(Ankudowich et al., 2016) and correct rejection reaction times (Cansino et al., 2012). These findings indicate that changes in age differences on context memory may occur around midlife.

Although the majority of studies aggregate males and females, studies that separate these groups have found notable sex differences in episodic memory (reviews by Andreano & Cahill, 2009; Asperholm et al., 2019; Herlitz & Rehnman, 2008). Females have a small and consistent advantage across episodic memory tasks than males (Andreano & Cahill, 2009; Asperholm et al., 2019; Herlitz & Rehnman, 2008). The magnitude of this advantage depends on stimulus material: Females perform better on verbal, nameable images, and location stimuli; Males perform better on spatial stimuli and abstract images. Crucially, sex differences in memory change over life. Cross-sectional evidence shows that females show more cross-sectional age differences on episodic memory than males in the sixties but not in the twenties or forties (Anstey et al., 2021), suggesting that the female memory advantage disappears after midlife. Similarly, longitudinal evidence shows substantial within-person episodic memory decline for women in their fifties (Karlamangla et al., 2017). A meta-analysis of 617 studies with over a million participants in total also established that sex differences may diminish with advancing age (Asperholm et al., 2019). Overall, these findings indicate that a female episodic memory advantage may diminish over adulthood, particularly at midlife.

Examining *when* changes occur in sex differences for aging of memory can offer key insights into the source of sex differences in memory. A change in the size or direction of sex differences that coincided with the occurrence of a major aging-related biological event could suggest that the biological factors were crucial in understanding the observed sex differences. Notably, the greater difficulties on episodic memory observed in some females after midlife (Anstey et al., 2021; Asperholm et al., 2019) occurs at the same time as one of the most significant endocrine events in a female’s life. Menopause typically occurs around ages 50 to 53 (Gold et al., 2013; Palacios et al., 2010), and marks the permanent cessation of menstruation as a result of loss in ovarian follicular function (Greendale et al., 1999). Menopause is characterized by a decline in sex hormone levels, particularly the ovarian hormone estrogen (Harlow et al., 2012), and this decline may lead to resulting memory decline as estrogen has neuroprotective effects and is a key modulator of learning and memory in the female brain (Galea et al,, 2017)..

Menopause has been associated with greater memory declines in females. Episodic memory declines in post-menopausal females compared to pre-menopausal females have been found in cross-sectional (Rentz et al., 2017; Weber et al., 2013; c.f., Greendale et al., 2009) and longitudinal studies (Epperson et al., 2013), and corroborated by a meta-analysis of existing evidence (Weber et al., 2014). A study of middle-aged adults found that pre- and peri-menopausal females had better memory than age-matched males on various memory tests including associative memory, but that post-menopausal females did not show a memory advantage (Rentz et al., 2017). Similarly, advanced age was associated with poorer context memory in post-menopausal females but not in pre-menopausal females (Crestol et al., 2023), suggesting age-related context memory decline emerges in late midlife after the menopausal transition. Taken together, these findings suggest that sex differences after menopause at midlife may be due to biological menopause-related factors and not chronological age.

The accelerated decline observed in episodic memory at midlife is interesting because it aligns with the time of menopause in midlife females, and menopause has been linked to greater memory decline. However, as most studies have not disaggregated aging of memory by biological sex, it is inconclusive whether the greater decline in memory is specific to females.

In the current study, we aimed to examine whether cross-sectional age differences in episodic memory differ for males and females. Importantly, disaggregating by sex enabled us to test whether past midlife effects on episodic memory for context were specifically driven by post-menopausal females, and whether males also showed decline at midlife. We predicted that both males and females would show little to no age-related decline until midlife, after which females would show accelerated decline coinciding with the timing of menopause.

We also investigated whether effects differed for context memory compared to item memory. In line with past findings, we predicted that the accelerated age-related decline at midlife would be specific to context memory, while item memory would remain relatively intact with age. In particular, females would show accelerated decline on context memory.

To quantitatively map aging, we use the statistical technique of changepoint regression (also called piecewise or segmented regression). This innovative technique is ideally suited for the current study as it quantifies parameters of theoretical interest: the rate of change per year of age (the slope, *β*), and the ages in which the rate shifts (the changepoint, *ψ*). Confidence intervals around the changepoint identify critical transition periods by estimating statistically significant changes in the regression line. Changepoint regression models are ideal for the current analyses because the changepoint has the theoretical utility of quantifying the timing when performance changes. Thus it mathematically quantifies the hypothesized theoretical process in which decline accelerates at midlife. This technique considerably advances past studies which have used linear regression, thus assuming a steady change in performance with age. It also advances past non-linear techniques such as polynomial regression or regression splines which either do not estimate changepoints or have values (e.g., maximum/minimum points) that are not intuitive to interpret.

Evidence on the influence of menopause on memory suggests that effects of chronological age may be due in part to reproductive aging. However, chronological aging and reproductive aging are correlated processes and hence are difficult to separate methodologically (Taylor et al., 2019). Age and menopause cannot be entered in the same statistical model as they share considerable variance. Our approach addresses this issue by modelling the effects of age alone, and then examining if estimated age changepoints align with the observed timing of menopause. By comparing age-matched males, we are also able to identify any sex differences in memory decline, which would indicate effects specific to menopausal endocrine aging rather than chronological aging. The innovative approach of examining sex differences using changepoint regression allows us to address the question of whether shifts in context memory decline with age may be due to reproductive-related processes rather than chronological aging.

## Research Design and Methods

### Participants

We used existing data from the Brain Health at Midlife and Menopause (BHAMM) study (Crestol et al, 2023). Participants were 192 adults aged 21 to 65 (M=44, SD=13), including 134 females with ovaries. Self-reported ethnicity data recorded 74.5% White, 6.3% East Asian, 4.7% Latin American, 3.6% South Asian, 3.1% Black, 3.1% Mixed, 2.6% West Asian, 1.0% South-east Asian, 0.5% North African, and 0.5% Indigenous. Tables 1 and 2 show the descriptive statistics for age and education by sex and menopause stage.

**Table 1.**
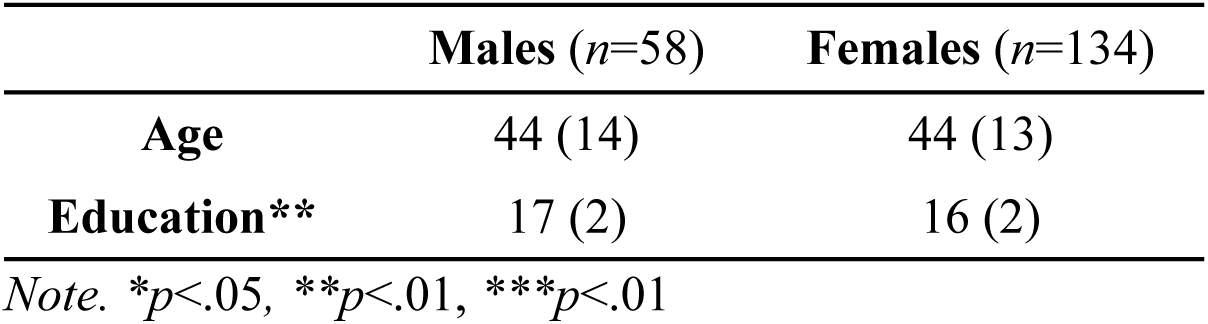
Mean (*SD*) for age and education per sex.

|  | **Males (*n*=58)** | **Females (*n*=134)** |
| --- | --- | --- |
| **Age** | 44 (14) | 44 (13) |
| **Education\*\*** | 17 (2) | 16 (2) |

*Note.* \* $p<.05$ , \*\* $p<.01$ , \*\*\* $p<.01$

**Table 2.** Mean (*SD*) for age and education per menopause stage.

|  | **Pre-menopause (*n*=85)** | **Post-menopause (*n*=49)** |
| --- | --- | --- |
| **Age\*\*\*** | 36 (10) | 57 (4) |
| **Education** | 16 (2) | 15 (2) |

*Note.* \* $p<.05$ , \*\* $p<.01$ , \*\*\* $p<.01$

Participants were recruited using advertisements in the Montreal community and over the internet. Participants signed an online consent form and completed a screening questionnaire with demographic information, medical history, and reproductive history. Inclusion criteria were a high school education or higher, willingness to provide a blood sample, and good mental and physicalhealth. Exclusion criteria were untreated cataract/glaucoma/age-related maculopathy, uncontrolled hypertension, untreated high cholesterol, diabetes, history of estrogen-related cancers, chemotherapy, neurological diseases, history of serious head injury, history of major psychiatric disorders, claustrophobia, history of a substance use disorder, currently smoking >40 cigarettes per day, a BMI>40, and inability to meet requirements for magnetic resonance imaging (MRI) safety (e.g., implanted pacemaker). Females were also excluded for current use of hormone replacement therapy, bilateral oophorectomy, current pregnancy, current use of hormonal birth control for a reason other than contraception, perimenopausal status, or indeterminate menopause status (based on self-report and hormonal data),

### Procedure

Participants who met the inclusion and exclusion criteria were invited to complete an initial testing session in which they completed an MRI safety questionnaire, a battery of neuropsychological and psychiatric assessments, donated blood samples for hormonal assays, and performed a practice version of the spatial context memory task in a mock MRI scanner. The neuropsychological assessments included the Mini International Neuropsychiatric Interview (M.I.N.I; Sheehan et al., 1998; exclusion criteria was indication of undiagnosed psychiatric illness), the Beck Depression Inventory II (BDI-II; Beck et al., 1997; inclusion cut-off ≤ 19), and the Mini Mental State Examination (MMSE; Folstein et al., 1975; inclusion cut-off ≥ 27). Participants who met the neuropsychological inclusion criteria, were classified as pre- or post-menopausal, and were able to perform the practice fMRI task were invited to a second testing session. In the second session, they completed a spatial context memory task while . functional magnetic resonance imaging (fMRI) scans were collected. Further details of hormonal staging, the fMRI session, and other materials and procedures are available in Crestol et al (2023). In the current study we focus on the behavioral fMRI data and neuropsychological data from the first visit.

## Measures

### Hormonal staging

Blood was drawn from non-fasting individuals by a certified research nurse during Sessions 1 and 2. Collecting blood draws in both sessions enables confirmation that hormonal staging has not changed between sessions, which is particularly relevant for midlife females. Levels of Estradiol-17β (E2), Follicle Stimulating Hormone (FSH), Luteinizing Hormone (LH), and Progesterone were assessed. Menopausal staging was based on menstrual cycling and the Stages of Reproductive Aging Workshop (STRAW) +10 criteria (Harlow et al., 2012). Participants were categorized as pre-menopausal or post-menopausal using self-report and measured hormonal levels for FSH using STRAW+10 criteria.

### Spatial Context Memory Task

E-Prime software (Psychology Software Tools, PA) was used to present a spatial context memory task, and to collect response accuracy and reaction times. Participant responses were collected using two 4-button inline fibre optic response boxes. During the task, response options and the corresponding buttons were displayed at the bottom of the screen to ensure clarity.

Task difficulty was manipulated by increasing encoding load. Participants encoded blocks of either six faces (an Easy condition) or twelve faces (a Hard condition), to enable calculation of participant performance as a function of increased task demands. Past studies using the same task have shown that age differences may interact with task difficulty (Ankudowich et al., 2016; c.f., Kwon et al., 2016), but our main measures of interest (Sex and Sex by Age) were not different when parsing by task difficulty, hence easy and hard conditions were averaged in the current analyses for simplicity.

The task consisted of encoding and retrieval phases, described below. At the end of each phase, participants were given 60 seconds to provide a confidence rating of their performance on a 4-point scale. The total duration of the task was approximately 1 hour, 7 minutes, and 44 seconds.

#### Encoding Phase

Instructions were presented for the first six seconds of each encoding phase. Participants were instructed that they would see a series of black and white photographs of faces, and told to memorize each face and its spatial location. To help ensure that participants deeply encoded the stimuli, they also completed a pleasantness rating (Bernstein et al., 2002) in which they rated each face as pleasant or neutral. Further details on the stimuli used are available in Rajah et al. (2008).

Participants were then presented with blocks of 6 (easy condition) or 12 (hard condition) face stimuli in sequence. Faces varied in age (from toddlers to older adults), biological sex, and ethnicity. Each face was presented one at a time for two seconds in one of four quadrants on the screen (upper right, upper left, lower right, lower left). The inter-trial interval (ITI) between face stimuli varied from 2.5 to 7.5 seconds (mean ITI=4.17 seconds).

Participants completed a minimum of two easy task runs in the encoding phase, consisting of two easy task blocks of six stimuli each (total duration 9:42 minutes). Hence each participant experienced at least 24 encoding events in the easy condition (4 task blocks x 6 stimuli). Participants completed a minimum of three hard task runs in the retrieval phase, consisting of one hard task block of 12 stimuli. Hence each participant experienced at least 36 encoding events in the hard condition (3 task blocks x 12 stimuli). Participants completed a maximum of four easy and hard task runs, however due to participation withdrawals and software errors, not all participants experienced all runs. The order of runs was counterbalanced across participants.

#### Retrieval Phase

Instructions were presented for 14 seconds. Participants were informed that they would see a series of previously encoded faces (old) and novel faces (new). They were told to respond to each face by pressing a button corresponding to one of six responses: identifying the face as new, familiar but they didn’t remember the location, or remembering the face and its location (one of the four quadrants). Participants were informed not to guess and to only respond with a location if they clearly recollected the face and its location.

Participants were then presented with blocks of 12 (easy condition) or 24 (hard condition) faces in sequence. Each face was presented one at a time for four seconds in the centre of the screen. The inter-trial interval (ITI) between face stimuli varied from 2.5 to 7.5 seconds (mean ITI=4.17 seconds). Participants viewed a total of 96 faces in the retrieval phase, 48 old and 48 new.

Participants completed a minimum of two easy task runs in the retrieval phase, consisting of two easy task blocks of 12 stimuli each. Hence each participant experienced at least 48 retrieval events in the easy condition (4 task blocks x 12 stimuli), 24 old and 24 new. Participants completed a minimum of three hard task runs in the retrieval phase, consisting of one hard task block of 24 stimuli. Hence each participant experienced at least 72 retrieval events in the hard condition (3 task blocks x 24 stimuli), 36 old and 36 new.

Participants were excluded from analyses if they did not provide at least one correct spatial context retrieval response (*n*=11), were below chance for correct rejections (*n*=1), or had a Cook’s D > 3 SD above the mean on 3 or more measures after regressing age on each measure (*n*=2). The final sample for the spatial context memory task was 178 participants.

#### Outcome measures

Accuracy (% correct) and reaction time (milliseconds) were calculated for each of the following response types.

1. <u>Correct Spatial Context Retrieval:</u> Total number of correct associative retrieval trials, calculated by the total responses correctly recalling previously encoded faces and locations divided by the total previously seen faces presented at retrieval. This measure indexes *spatial context memory*.
2. <u>Correct Recognition:</u> Total number of *correct retrieval* trials, calculated by the total responses correctly recognizing previously encoded faces without location divided by the total previously seen faces presented at retrieval. This measure indexes *facial item memory*.

### Verbal Memory Task

Verbal memory was assessed using the California Verbal Learning Test (CVLT-II; Delis et al., 2000). English and French versions of the task were used. Participants were excluded from analyses if their first language was not English or French (*n*=32). The final sample for the spatial context memory task was 161 participants (113 females).

Participants were auditorily presented with two 16-item word lists: a test list and an interference list. The first (test) list consists of four words from four semantic semantic categories verbally presented five times (i.e., over five trials), with participants asked to recall the words after each trial. The second (interference) list consists of 16 distractor words verbally presented one time (i.e., for one trial). Free and semantically-cued recall of the first list occurs immediately after the second list (short recall), and again after a 20-minute delay in which non-verbal material is presented (long recall). Finally, participants complete a recognition memory test in which they indicate (via “yes” or “no”) whether they recognize words from the test list mixed with words not from the test list. Further details on task stimuli and methodology are available in Delis et al (1987, 2000, 2001).

#### Outcome measures

The number of correct responses was collected for the following response types and converted to accuracy (% correct). All three measures index *verbal item memory*.

1. <u>Long Free Recall:</u> Total number of correctly learned items (from the test list) recalled with no cue divided by the total number of learned items.
2. <u>Long Cued Recall:</u> Total number of correctly learned items (from the test list) recalled with a semantic cue (one cue for each of the four semantic categories) divided by the total number of learned items.
3. <u>Recognition:</u> Total number of correctly recognized items (from the test list) divided by the total number of learned items.

### Analyses

Analyses were conducted with version 4.3.0 of the R language and environment for statistical computing (R Core Team, 2023), with the *segmented* package version 1.6.4 to run the Davies test and changepoint regression (Muggeo, 2008).

### Changepoint regression to estimate non-linear relationships

Analyses were separately conducted for each response measure and sex. To first establish if there was a significant non-linear relationship, we used a Davies test (Davies, 2002) for a non-zero difference in the slope parameter. Next, non-linear age effects were estimated using changepoint regression. Changepoint regression was conducted with a linear reparameterization approach which iteratively fits piecewise terms in a regression model (Muggeo, 2003, 2008). Changepoint regression models included terms for the changepoints (*ψ*) and the slopes (*β*) before and after the changepoint. A Welch two-sample t-test revealed that education differed significantly between the sexes, *t*(126) = 2.6, *p* = .01, 95% CI 0.2, 1.5, hence education was included as a covariate in the regression models to control for confounding effects of education on episodic memory.

A hierarchical regression procedure was used in which increasingly complex models were fitted, beginning with a linear model [1] with constant effects of age and education on episodic memory,

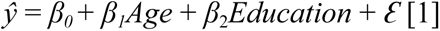

followed by a one-changepoint model [2] in which the linear effect of age was allowed to shift at any one location,

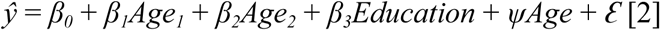

and lastly a two-changepoint model [3] in which the linear effect of age was allowed to shift at any two locations.

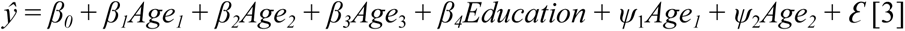

Increasingly complex models were compared inferentially with a chi-squared difference test and on model parsimony using the Akaike information criterion (AIC). A best-fitting model was selected if it offered a significantly better fit than a simpler model (*p*<.05) and an improvement in parsimony (ΔAIC > -2).

Estimates of the parameters of interest are presented for the final selected model using raw effect sizes. Raw effect sizes use the original units of measurement (i.e. percentage of correct responses for accuracy and milliseconds for reaction time) and hence are intuitive to interpret. The slope coefficients (*β*) estimate the difference in performance per year of age and the changepoint values (*ψ*) estimate the age(s) in which performance shifts.

## Results

### Spatial Context Memory Task

#### Correct Spatial Context Retrieval

Results are shown in Table 3. On accuracy for males, the best fitting model was a linear model with a non-significant slope (*β=*-2.2, *SE*=2.1, *t*=-1.1, *p*>.05), indicating that correct spatial context retrieval did not vary significantly from young adulthood to the end of mid-adulthood. On accuracy for females, the best fitting model was a one-changepoint model, in which accuracy significantly declined gradually by 2% per year of age from young adulthood until mid-adulthood (*β_1_=*-2.1, *SE*=1.0, *t*=-2.0, *p*<.05), shifting at age 49.6 (95% CI 41.2, 58.1) to more rapid decline of 4% per year of age until the end of mid-adulthood (*β_2_=*-3.7, *SE*=1.1, *t*=-3.3, *p*<.05). On response times the best fitting models were linear for both sexes, with no significant differences in response times across ages for males (*β=*3.6, *SE*=21.4, *t*=0.1, *p*>.05) and females (*β=*34.5, *SE*=27.0, *t*=1.3, *p*>.05).

**Table 3.**
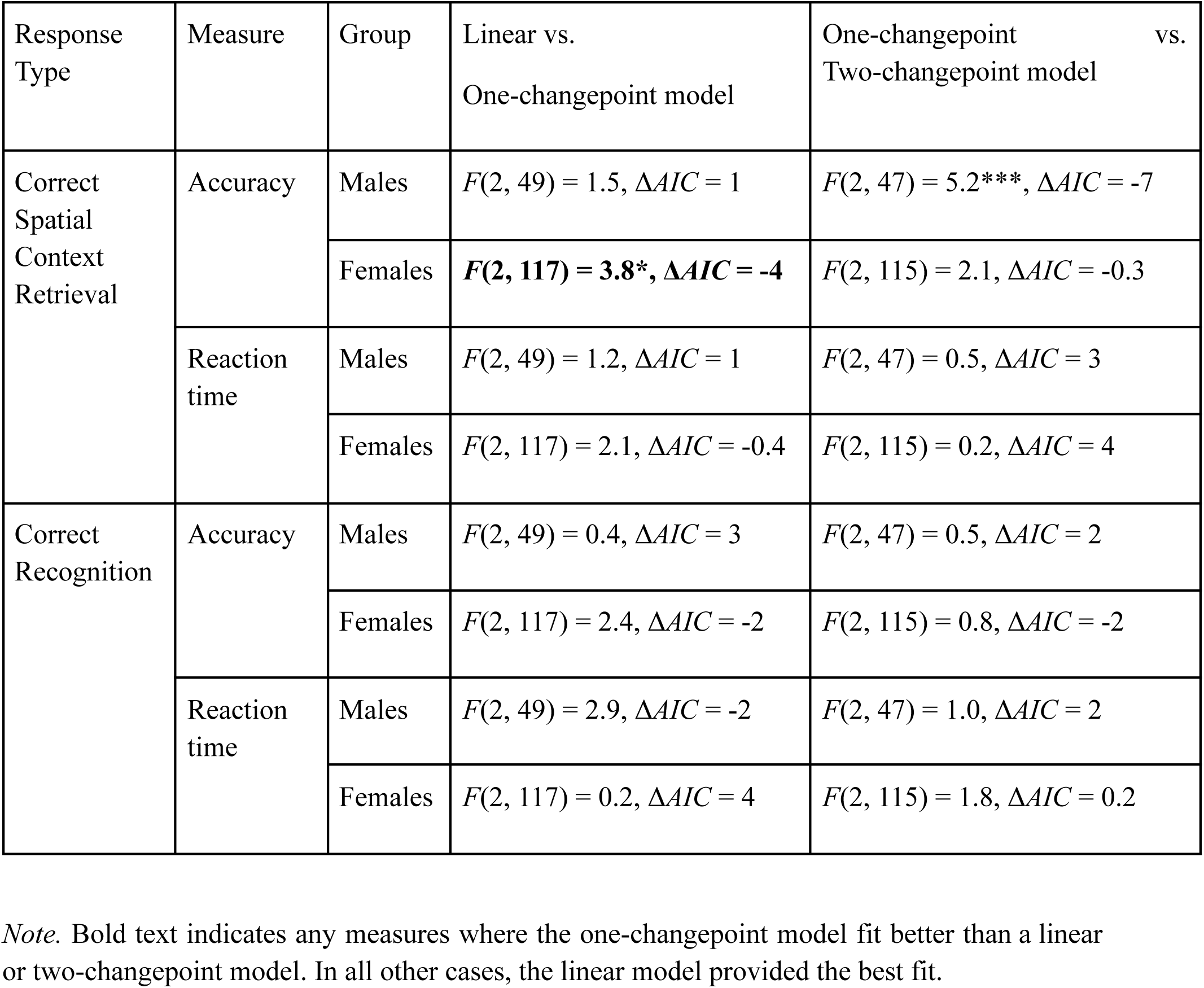
Hierarchical Regression Results Comparing Fitted Models on the Spatial Context Memory Task.

**Table 4.** Hierarchical Regression Results Comparing Fitted Models on the Verbal Memory Task.

| Response Type | Group | Linear vs. One-changepoint model |
| --- | --- | --- |
| Long Free Recall | Males | $F(2, 40) = 0.6, \Delta AIC = 3$ |
| | Females | $F(2, 99) = 1.6, \Delta AIC = 1$ |
| Long Cued Recall | Males | $F(2, 40) = 1.2, \Delta AIC = 0.2$ |
| | Females | $F(2, 99) = 2.0, \Delta AIC = -0.2$ |
| Recognition | Males | $F(2, 40) = 1.2, \Delta AIC = 1$ |
| | Females | $F(2, 99) = 1.7, \Delta AIC = 1$ |

*Note.* In all cases the linear model provided the best fit.

#### Correct Recognition

On accuracy and response times, the best fitting models were linear for both sexes. Males had no significant differences in correct recognition responses across ages (*β=*-0.1, *SE*=1.7, *t*=-.1, *p*>.05). Females had 2% more correct recognition responses across ages (*β=*2.0, *SE*=0.8, *t*=2.6, *p*<.05). Both males (*β=*-45.1, *SE*=88.6, *t*=-0.5, *p*>.05) and females (*β=*-30.9, *SE*=37.2, *t*=-0.8, *p*>.05) had no significant differences in response times across ages.

### Verbal Memory Task

There were no significant effects of age nor sex on the CVLT long free recall for males (*β=-*0.1, *SE*=2.0, *t*=-0.04, *p*>.05) and females (*β=*0.5, *SE*=0.9, *t*=0.5, *p*>.05) long cued recall for males (*β=*-1.8, *SE*=1.8, *t*=-1.0, *p*>.05) and females (*β=-*0.02, *SE*=0.9, *t*=-0.03, *p*>.05) long delay recognition for males (*β=*-0.7, *SE*=0.9, *t*=-0.7, *p*>.05) and females (*β=*0.2, *SE*=0.4, *t*=0.5, *p*>.05).

## Discussion and Implications

The current findings reveal that cross-sectional age differences in episodic memory for spatial context differ by sex. Males and females show similar age differences from young adulthood until midlife on spatial context memory accuracy, after which females show greater age differences than males. Using changepoint regression we quantified the critical transition period in which sex differences emerged as being around 50 years of age . Importantly, the timing when females show accelerated decline in context memory aligns with the timing of menopause in midlife females, which occurs around 50 years of age both in existing research on North American females (Gold et al., 2013; Palacios et al., 2010) and in the current sample.

Past studies have largely demonstrated independent effects of chronological age or reproductive age (menopause status) on episodic memory, as well as sex differences in the effect of chronological age on memory. However, few studies have examined these effects together, partly because of the methodological challenge of overlapping variance between age and menopause (Taylor et al., 2019). Our approach addresses this challenge by modelling age differences independently, within sex, and then examining if shifts in the timing of age differences align with the observed timing of menopause.

We found accelerated decline in females but not males specifically on spatial context memory (i.e., correctly recalling the source/location of previously learned information) and not on facial recognition or verbal item memory (i.e., correctly recognizing or recalling previously learned information). Our findings align with past work that age differences in context memory are particularly pronounced on binding memory for context rather than on memory for content (Chalfonte & Johnson, 1996). The results support the associative deficit hypothesis of a greater difficulty with advanced age specifically on binding pieces of information (Old & Naveh-Benjamin, 2008; Naveh-Benjamin, 2000). Our results also extend these findings to show that difficulties with advanced age on recalling the context or binding/associating items of learned information also differs critically by biological sex.

A strength of this study is using a sample of individuals that have been carefully and systematically categorized into menopause stages by using endocrine data collected with blood draws and published staging criteria. Thus effects of chronological age in females could be examined alongside reproductive age. By comparing models fitted in females and males of the same ages, we could examine whether episodic memory changes were specific to sex-specific factors rather than chronological aging. The current findings highlight the importance of studying middle-aged samples, because sex differences in episodic memory emerged at midlife and aligned with a midlife-specific reproductive event.

The findings also highlight the importance of studying sex differences in cognitive aging. The majority of existing cognitive aging studies have aggregated males and females, thus assuming no sex differences in cognitive performance with age. The current findings align with emerging evidence that biological sex is a critical factor when studying lifespan trajectories (Jacobs & Goldstein, 2018; Taylor et al., 2019). In additional to typical aging, it is valuable to elucidate sex differences in episodic memory, because larger episodic memory deficits than in normal aging are a hallmark of Alzheimer’s disease (Gallagher & Koh, 2011), and females are at greater risk of developing Alzheimer’s disease than males (Alzheimer’s Association, 2023; Andrew & Tierney, 2018; Podcasy & Epperson, 2016).

We observed linear decline in response times in both sexes. Our findings align with past work which shows that reaction times on context memory tasks are also slower by midlife (Ankudowich et al., 2012; Cansino et al., 2012). However, we did not observe any sex differences on response times across ages.

We observed an effect of task modality on sex differences in episodic memory, with non-linear decline and sex differences on spatial context memory recall, compared to linear decline with no sex differences on verbal memory recognition/recall and on facial memory recognition. Past work has shown that females have an advantage over males on tasks with verbal or facial information (Bender et al., 2010; Bleecker et al 1988; Erngrund et al., 1996a, 1996b; van Hooren et al. 2007), while males have an advantage on tasks with spatial information (Caselli et al., 2015; Gallagher & Burke, 2007; Lewin et al., 2001). It could be that the female advantage on verbal/facial information diminishes sex differences in aging on tasks with such information, while sex differences are more pronounced on spatial tasks. However, other studies have found non-linear decline with sex differences on episodic memory tasks with verbal stimuli, but linear decline on other stimuli such as faces and locations (Asperholm et al., 2019). Hence before conclusions can be drawn on sex differences in task modality, further research is needed to isolate effects of task modality from task design. As our spatial and verbal memory tasks differed both in task design and modality, future studies can use multiple tasks per modality, or use tasks matched on design elements but different on modality (although it is challenging to develop a task that measures context with verbal or facial stimuli). One design difference between our tasks is that the verbal task (CVLT) involves a short list of items with repeated verbal presentation, while the spatial task involves a longer number of items without repeated presentation. Thus the tasks have differing difficulty levels and also measure item memory differently. As age effects may be influenced by task difficulty (e.g., Ankudowich et al., 2016) it is possible that the more difficult spatial memory task was more sensitive to detect aging.

A methodological limitation of the current study is using a cross-sectional sample to examine chronological age differences. Future work with a longitudinal sample is needed to confirm that observed age differences index age-related decline. A second methodological limitation is the smaller sample size for males compared to females.

Future research can examine the stage of perimenopause more closely. We did not include perimenopause in the current analyses as peri-menopause is methodologically challenging to study: It is highly variable compared to pre- and post-menopause, and we could not distinguish between short- and long-term effects on cognition. Cognitive function does not change linearly across perimenopause (Greendale et al., 2009; Weber et al., 2013), and some short-term effects on cognition are reversed (Weber et al., 2013). Hence a peri-menopause group is statistically different from pre- and post-menopause groups due to the heterogeneity of variance, and is theoretically different due to the combination of both short- and long-term cognitive influences.

In conclusion, we observed critical sex differences in the aging of episodic memory for spatial context over adulthood. Our findings show that examining variability in the timing of sex differences over adulthood can offer key insights into the source of sex differences. Our findings suggest that menopause-related biological changes may underlie observed sex differences in memory decline. By comparing to age-matched males, we were able to establish that greater spatial context memory decline at midlife was specific to menopausal endocrine aging rather than chronological aging.

**Figure 1.**
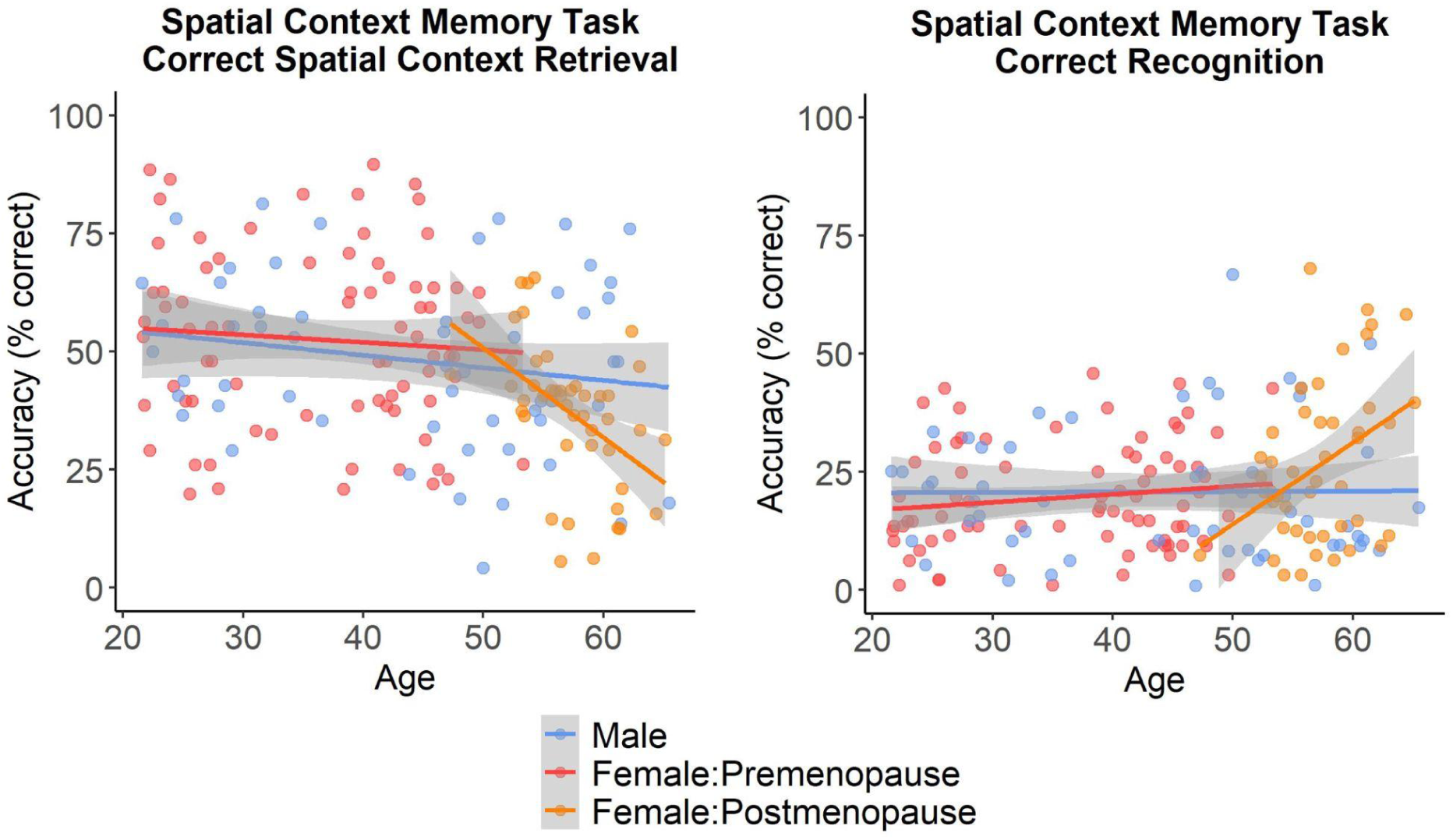
Scatterplot of Performance on the Spatial Context Memory Task Accuracy per Sex and Menopause Group. *Note*. Dots indicate individual participant performance, and lines indicate a line-of-best fit for each sex and menopause group.

**Figure 2.**
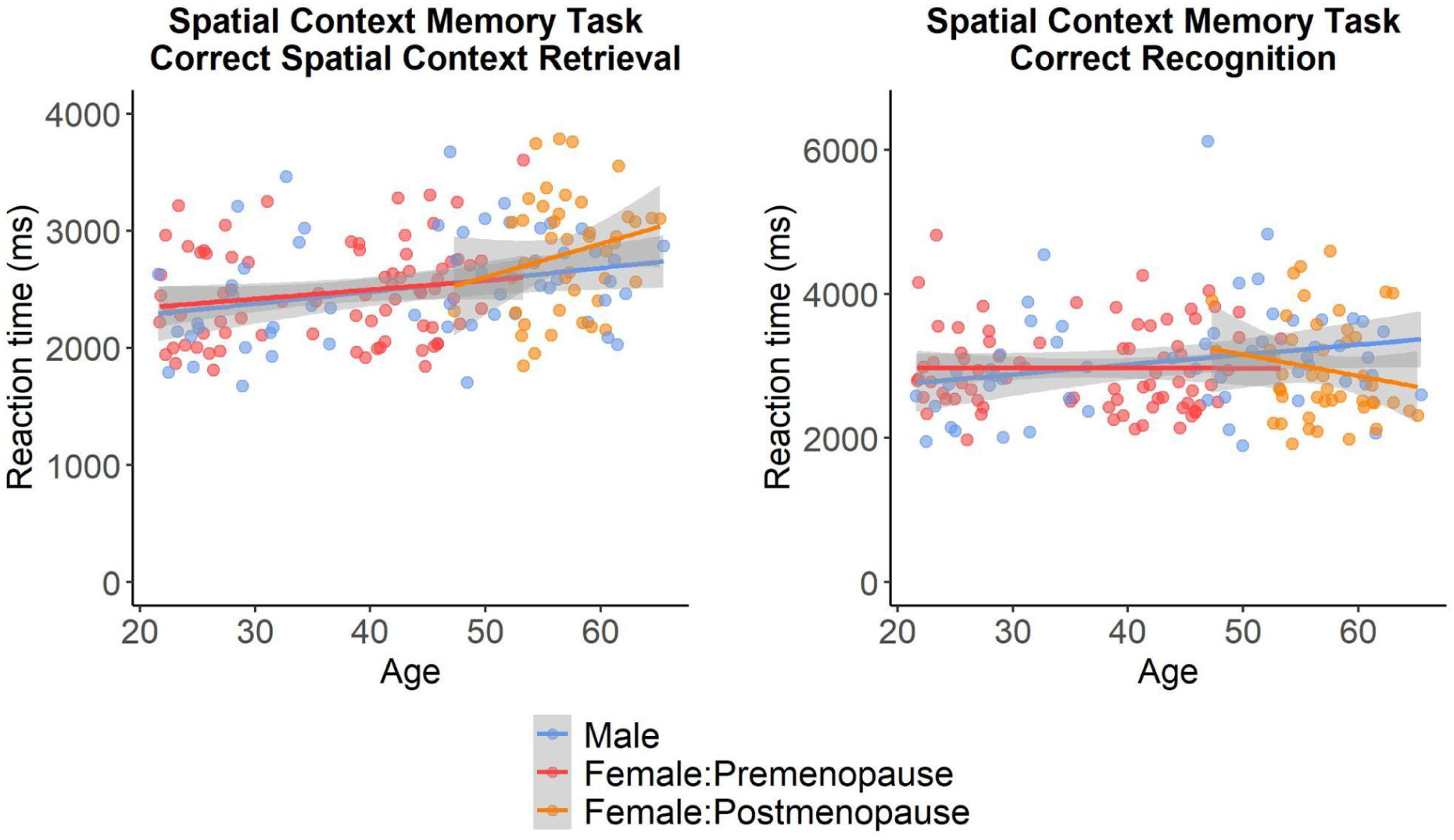
Scatterplot of Performance on the Spatial Context Memory Task Accuracy per Sex and Menopause Group. *Note*. Dots indicate individual participant performance, and lines indicate a line-of-best fit for each sex and menopause group.

**Figure 3.**
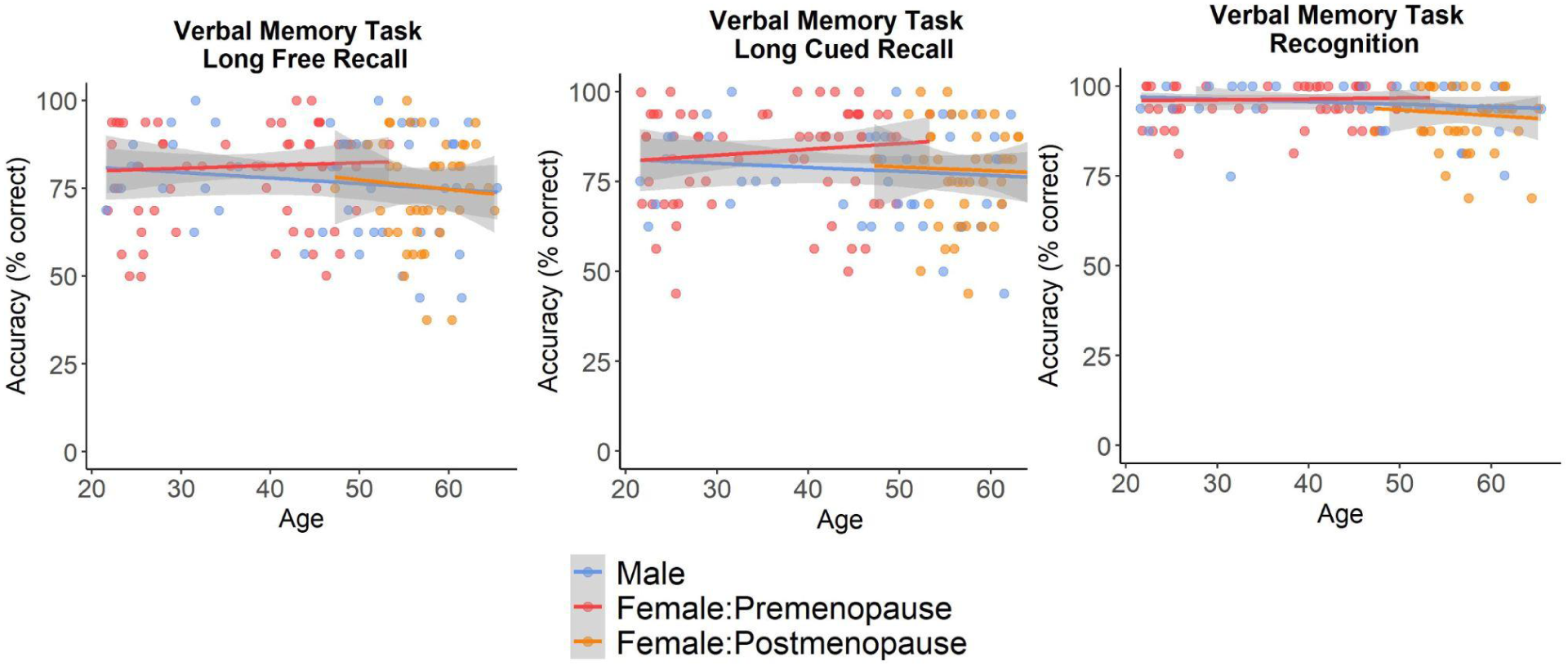
Scatterplot of Performance on the Verbal Memory Task per Sex and Menopause Group. *Note*. Dots indicate individual participant performance, and lines indicate a line-of-best fit for each sex and menopause group.

## Acknowledgements

We thank the research participants of the Brain Health at Midlife and Menopause (BHAMM) Study for their valuable contributions to this project. We thank S. Pasvanis for her work on database administration and coordination. We also thank the part-time research assistants (H. Azizi, R. Young, A. Condescu, L.Khyatt) and trainees (S. Subramaniapillai, G. Velez Largo, J. Kearley, A. Duval, J. Snytte, S. Loparco) who helped with participant recruitment, testing, or MRI data quality control. We are grateful for the support of the Canadian Consortium on Neurodegeneration in Aging (CCNA) Teams 9, 10, and WSGD Theme, the MRI Staff at the CIC, DMHUI, and Dr. D. Cohen for help with recruitment.

## CREDIT Authorship Contributions

AAL: Conceptualization, Methodology, Software, Formal analysis, Writing - Original Draft, Writing - Review & Editing, Visualization, Funding acquisition; RL: Formal analysis, Validation JK: Writing - Review and Editing; MNR: Conceptualization, Methodology, Resources, Writing - Review & Editing, Supervision, Project administration, Funding acquisition

